# 4’-fluorouridine is a potent inhibitor of Oropouche virus in vitro and in animal infection models

**DOI:** 10.1101/2025.09.22.677733

**Authors:** Martin Ferrié, Maïlis Darmuzey, Itona Tarillon, Thibault Tubiana, Mona Khan, Tania Roskams, Birgit Weynand, Dietmar Rudolf Thal, Niels Cremers, Stijn Hendrickx, Kim Donckers, Thayara Morais Portal, Bert Vanmechelen, Viktor Lemmens, Joana Rocha-Pereira, Concetta Castilletti, Peter Mombaerts, Stéphane Bressanelli, Manon Laporte, Hélène Malet, Johan Neyts

## Abstract

Oropouche virus (OROV) is an orthobunyavirus that causes increasingly frequent and severe outbreaks in Central and South America. We report that 4′-fluorouridine (4′-FlU) inhibits the *in vitro* replication of epidemic and pre-epidemic OROV strains in multiple cell lines. *In vitro* polymerase assays demonstrate that 4’-FlU (as its triphosphate) targets the *Peribunyaviridae* L protein, is incorporated during RNA synthesis and causes premature chain termination. Following 69 consecutive days of *in vitro* passages of OROV in the presence of suboptimal concentrations of 4’-FlU, no drug-resistant variants were identified in the viral polymerase. In stringent mouse (AG129) or Syrian hamster OROV-infection models, oral administration of 4′-FlU completely blocked viral replication and virus-induced disease, even when administration was delayed until 72 hours after infection. Our findings support exploring the potential of 4’-FlU for the management of OROV infections in humans.

## INTRODUCTION

The Oropouche virus (OROV, *Orthobunyavirus* genus, *Peribunyaviridae* family)^1^ is an arthropod-borne virus, transmitted mainly by the midge *Culicoides paraensis*. It was first isolated from humans in 1955 in the Oropouche region of the Caribbean island Trinidad^2^. OROV has emerged as a significant arboviral threat across Central/South America causing recurrent outbreaks, with cumulative case numbers estimated in the hundreds of thousands^3^. Strikingly, according to the Pan American Health Organization (PAHO), Brazil experienced a sharp rise in Oropouche fever cases, with 11,888 laboratory-confirmed cases in 2025, representing a consequential increase from the 831 cases reported in 2023^4^. Recently, OROV infections have soared in regions where the virus was not previously detected, including in Cuba and French Guyana^4^. The virus was also detected in non-endemic regions, including North America and Europe, primarily among travelers returning from endemic areas^3,5–7^. This increased detection of human cases may also be attributable to enhanced surveillance efforts and improved diagnostic capabilities, particularly outside the Amazon basin. OROV-associated symptoms in humans are defined as a febrile illness and are often mimicking dengue-like or Zika-like syndromes, making it difficult to assess the real burden of OROV^8,9^.

Unlike previous outbreaks, which involved only asymptomatic or mild cases of OROV infection in humans, the 2024 outbreak in Central and South America, marked the first time that human fatalities were reported, with two deaths linked to the virus^10^. In July 2024, the Pan American Health Organization issued an epidemiological alert about the potential of vertical transmission of OROV^11^. In December 2024, Brazil reported three confirmed cases of vertical OROV transmission, resulting in stillbirths or congenital malformation^12–14^. Given the swift geographic expansion of OROV, its potential to cause severe clinical outcomes (deaths and vertical transmission) and the toll such an epidemic can impose on healthcare systems, a strong and integrated global response, including medical countermeasures, is urgently needed^15^.

The OROV genome consists of three segments of negative-sense RNA, called Small (S; encoding the nucleocapsid N and non-structural Ns protein), Medium (M; encoding the surface glycoproteins Gn/Gc and the non-structural protein NSm) and Large (L; encoding the L protein that comprises the RNA-dependent RNA-polymerase)^3^. The segmented nature of the OROV genome enables reassortment events, a process commonly observed among segmented viruses^16^. Reassortment events have played a critical role in the emergence of current epidemic strains, resulting from the reassortment of the M segment from virus circulating between 2009 and 2018 in the eastern Amazon, and the L and S segments from viruses identified between 2008 and 2021 in Peru, Colombia and Ecuador^9,17,18^. These recent genetic exchanges, between different viral lineages, are considered a driving force behind the recent outbreaks and the expanding geographic range of OROV.

Despite the geographical expansion of OROV and its increased virulence in humans there are no antiviral drugs for the management of OROV infections. Treatment remains supportive, and is limited to relieving symptoms and preventing complications. The broader-spectrum RNA virus inhibitor ribavirin lacks antiviral activity against OROV^19^. Human and mouse interferon-α are active *in vitro* and *in vivo* (prophylactic dosing), respectively, but lose their protective effects in a mouse model, when treatment is initiated after infection^20^. This underscores the pressing need for an effective and safe drug for the treatment of OROV infection. We therefore assembled a panel of antiviral drugs and assessed their potential effect on OROV replication. This consisted in molecules with known broader-spectrum activity against a number of RNA viruses and (ii) baloxavir, an influenza endonuclease inhibitor that exerts modest *in vitro* antiviral activity against Bunyamwera virus (another *Orthobunyavirus*)^21^. The activity was assessed against both of an epidemic clinical OROV isolate (OROV-IRCCS)^22^ and a pre-epidemic reference OROV strain isolated in the 1960’s (OROV-TR9760). Only 4’-FlU, a molecule reported to exert antiviral activity against a number of viruses such as respiratory syncytial virus (RSV), Influenza A virus (IAV), severe acute respiratory syndrome coronavirus 2 (SARS-CoV-2), Lassa virus (LASV), Nipah virus (NiV) and chikungunya virus (CHIKV)^23–26^, was found to markedly inhibit OROV replication. These results prompted us to study its efficacy in animal infection models for which we used the 2024 epidemic clinical isolate OROV-IRCCS. We demonstrate that oral administration of 4’-FlU completely blocked viral replication and conferred protection against virus-induced disease in mice and hamsters.

## RESULTS

### 4’-FlU is a potent inhibitor of OROV replication in cell culture

We explored whether (i) a panel of molecules with reported antiviral activity against multiple RNA viruses, GS-441524 (the parent nucleoside of remdesivir), obeldesivir (an oral prodrug form of GS-441524), favipiravir, ribavirin, molnupiravir, galidesivir, sofosbuvir and 4′-Fluorouridine (4′FlU)], as well as (ii) baloxavir (an influenza endonuclease inhibitor with modest antiviral activity against Bunyamwera virus^21^), can inhibit the *in vitro* replication of an epidemic clinical isolate (OROV-IRCCS; obtained from a patient who returned from Cuba to Italy in July 2024)^22^ and a pre-epidemic reference OROV strain (isolated in the 1960’s; OROV-TR9760) in VeroE6, A549 and JEG3 cells (Supplementary Fig. 1). None of the molecules, except for 4’FlU, resulted in a notable antiviral effect. 4’-FlU inhibited OROV-induced cytopathic effects (CPE) formation [EC₅₀ values ranging from 0.012 µM (VeroE6) to 0.048 µM (JEG3)] and reduced viral RNA levels in culture supernatant [EC₉₀ values of 0.17 µM (A549) and 0.24 µM (JEG-3), Fig. 1a and b]. No detectable cytotoxicity was noted at the highest concentration tested in the antiviral assays (50 µM) (Fig. 1a).

**Figure 1.**
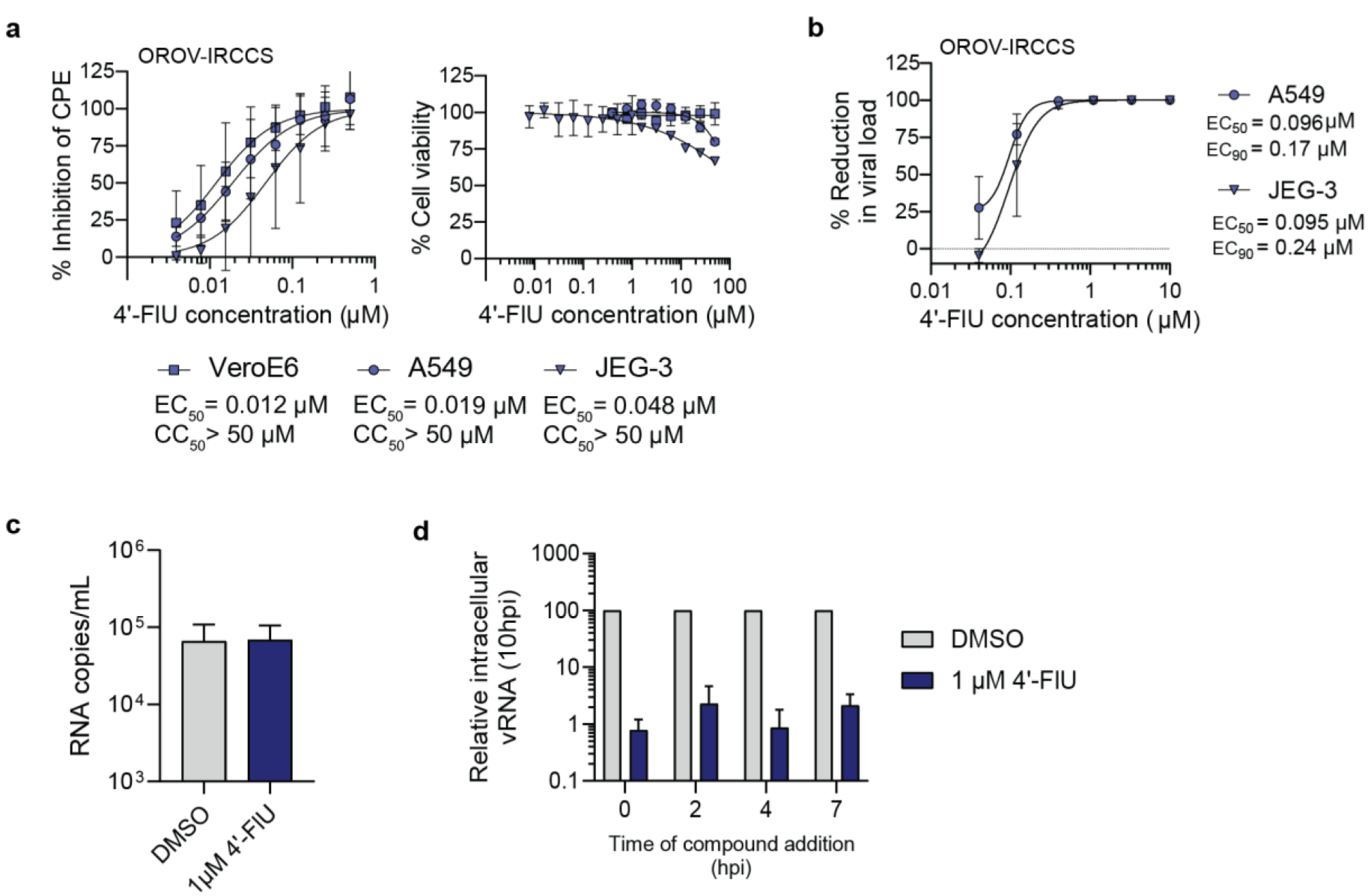
Antiviral activity of 4’ FlU against OROV in cell culture. **(a)** 4’FlU inhibits the replication of OROV in three cell lines. Dose–response curves of the antiviral activity of 4’-FlU determined by MTS-based CPE reduction assay. Cytotoxicity was determined by MTS cell viability assay in mock-infected cells. Data points are the mean ± SD of *n* = 4 biologically independent experiments. **(b)** OROV viral load reduction in A549 and JEG-3 cells based on the quantification of viral RNA in the supernatant by RT-qPCR 3 dpi. Data points are mean ± SD of n=3 biologically independent experiments. **(c)** Virucidal activity of 4’-FlU at 1µM against OROV in A549 cells, based on the quantification of viral RNA in the supernatant by RT-qPCR 1dpi. Data points are mean ± SD of n=3 biologically independent experiments. **(d)** Time-of-addition assay to determine at which stage of OROV replication cycle 4’-FlU (1 µM) is acting, based on the quantification of intracellular viral RNA by RT-qPCR 10hpi. Compound has been added to the cells at timepoints indicated on the x axis. Data points are mean ± SD of n=3 biologically independent experiments.

We next sought to dissect the mechanism of action of 4’-FlU. First, virucidal assays were performed in A549 cells to rule out any direct effect of the molecule on viral particles (Fig. 1c). At 1 µM, 4’-FlU had no effect on OROV-IRCCS infectivity, indicating that 4’-FlU mediates its activity exclusively intracellularly. We next performed time-of-addition assays in A549 cells (Fig. 1d). At 1 µM, 4’-FlU retained robust antiviral activity (> 97% inhibition of intracellular viral RNA loads), regardless of the time-of-addition following infection, supporting a post-entry mechanism of antiviral activity.

### 4′-FlU-TP acts as a chain terminator of *Peribunyaviridae* replication

As several reports highlighted that 4’-FlU target the polymerase of negative-sense RNA viruses and induce chain termination^25,26^, we here aimed to characterize whether 4′-FIU also targets the polymerase and interferes with the replication activity of the *Peribunyaviridae* L protein. Given that purified OROV L protein is not available, we performed *in vitro* replication assays with La Crosse (LACV) L protein (Fig. 2). LACV and OROV both belong to the *Peribunyaviridae* family, with LACV being a model virus for this family. The L protein of both viruses share 53.18% identity at the amino acid level (Supplementary Table 1). Importantly, 4’-FlU also efficiently inhibit the in vitro replication of LACV in VeroE6 cells (EC_50_ = 0.25 µM; Supplementary Fig. 3). The H34K mutant of the LACV L protein (LACV L_H34K_), which inactivates LACV L endonuclease activity^27^, was used for replication assays in order to prevent unwanted *in vitro* degradation of RNA by the LACV L endonuclease. LACV L_H34K_ was recombinantly expressed in insect cells, purified^28^, and successively incubated with (i) a mutated 5′vRNA containing the 5′ nucleotides 1-19 of the vRNA M segment (5′mut_1-19_) and (ii) a 25-mer 3′vRNA template containing the 3′ nucleotides 1 to 25 of the M segment (3′vRNA_1-25_). The 5′mut_1-19_ was used instead of wild type 5′vRNA to prevent *in vitro* formation of a double-stranded RNA between the highly complementary 5′ and 3′ vRNA, which would prevent *in vitro* replication activity^29^. The 5′mut_1-19_ had previously been shown to bind to LACV L in the same conformation as wild type 5′vRNA, and to promote correct binding of the 3′vRNA towards the LACV L active site, compatible with replication initiation^29^.

**Figure 2.**
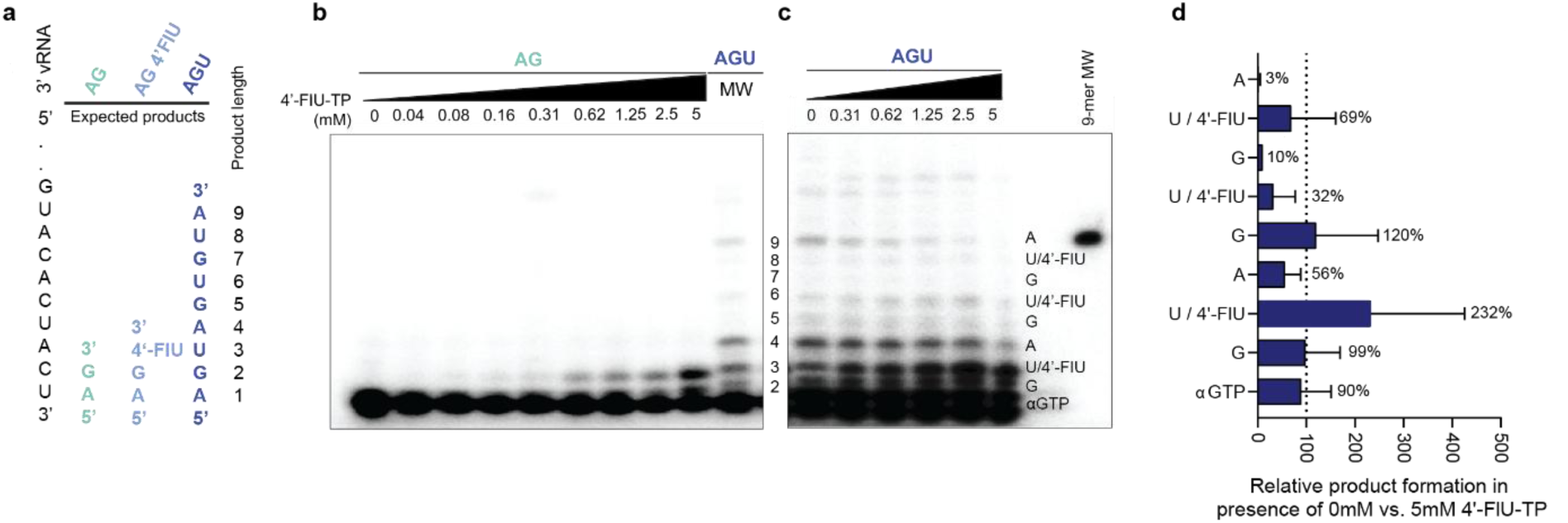
4′-FlU-TP acts as a chain terminator of LACV L_H34K_ replication *in vitro*. **(a)** Schematic representation of the 3′vRNA template and the expected replication products in presence of 2 NTPs (AG), 2 NTPs+4′-FlU-TP (AG 4′-FlU), or 3 NTPs (AGU). The expected product if 4′-FlU-TP acts as a chain terminator is shown for the AG 4′-FlU condition. Product length is indicated on the right. (**b)** LACV L_H34K_ *in vitro* replication assays in presence of 5′mut_1-19_, 3′vRNA_1-25_, 2 NTPs (AG) or 3 NTPs (AGU), MgCl_2_, and [^32^P-GTP]. 4′-FlU-TP is added at different concentrations and reactions are incubated for 4 h at 30 °C. The proposed length of the products is indicated on the right of the gel. A representative gel from 3 biologically independent experiments is shown. (**c)** LACV L_H34K_ *in vitro* replication assays with a competition between 100 μM UTP and increasing concentrations of 4′-FlU-TP. The molecular marker (MW) is a commercial RNA corresponding to a radiolabeled 9-nucleotide product. Representative gel from 3 biologically independent experiments is shown. **(d)** Histogram showing the ratio of replication products from reactions with 5 mM 4′-FlU-TP versus 0mM 4′-FlU-TP, calculated from Fig. 2b, shown as a function of product size. The proposed nature of the last nucleotide incorporated to the product is indicated. Data points are mean ± SD of n=3 biologically independent experiments.

*De novo* replication assays were performed in presence of MgCl_2_, ATP, GTP and increasing concentrations of 4′FlU-TP. A reaction in presence of LACV L_H34K,_ 5′mut_1-19,_ 3′vRNA_1-25,_ MgCl_2_ and the three-nucleotide subset containing ATP, GTP and UTP shows, using Urea-PAGE gels, the formation of the expected nine-nucleotide product (Fig. 2a-b)^29^. Analysis of the products formed when UTP is replaced with 4′FlU-TP indicates that incubation with increasing concentrations of 4′FlU-TP leads to an increased detection of a product interpreted to correspond to a tri-nucleotide product (Fig. 2a-b). These results suggest that 4′FlU-TP is incorporated but prevents product elongation (Fig. 2b). Competition experiments performed in presence of 100μM UTP and varying concentrations of 4′FlU-TP reveal that the 9-mer product gradually disappears when increasing concentration of 4′FlU-TP are added (Fig. 2c). Only 3% of the 9-mer product was formed in presence of 5 mM 4′FlU-TP compared to the same reaction performed in the absence of 4′FlU-TP (Fig. 2d). Consistently, increasing amounts of the 3-mer product are formed when increasing amounts of 4′FlU-TP are added, with a 232% increase in presence of 5 mM 4′FlU-TP compared to 0mM control (Fig. 2c and 2d). Collectively, these data show that 4′FlU-TP interferes with the replication activity of the *Peribunyaviridae* L protein by being incorporated in the reaction product, thereby acting as a chain terminator in *in vitro* replication assays.

### An OROV-IRCCS infection model in AG129 mice

To assess whether the potent *in vitro* antiviral activity of 4’-FlU translates to efficacy in an OROV animal infection model, we first aimed to establish such a model using the epidemic strain OROV-IRCCS (Supplementary Fig. 3). AG129 mice (which lack functional type I and type II IFN receptors) are known to be susceptible to infection with pre-2024 epidemic OROV strains^30^. In a pilot study, AG129 mice were subcutaneously inoculated in the footpad with 10⁵ TCID₅₀ of OROV-IRCCS.

Infected mice developed a severe acute disease characterized by reduced activity, weakness, progressive weight loss and a decline in surface body temperature. Clinical deterioration progressed quickly, with some mice reaching humane endpoints (HEP) as early as 3 days post-infection (dpi) and a cumulative lethality of 80% at day 7, the study endpoint (Supplementary Fig. 3b-e). At euthanasia, viral RNA and infectious virus titers were quantified in serum, brain, lungs, liver and spleen (Supplementary Fig. 3f-g). High levels of viral RNA were detected in infected mice [mean of 2.16×10^10^ RNA copies/mL serum; 3.91×10^8^ RNA copies/g brain; 2.10×10^10^ RNA copies/g lung, 4.59×10^10^ RNA copies/g liver and 7.49×10^9^ RNA copies/g spleen] (Supplementary Fig. 3f). Likewise, infectious titers were very high [4.05×10^8^ TCID_50_/mL serum, 2.32×10^6^ TCID_50_/g brain; 4.47×10^7^ TCID50/g lung; 4.41×10^9^ TCID_50_/g liver and 1.06×10^9^ TCID50/g spleen] (Supplementary Fig. 3g). Given the high level of viral replication, AG129 mice can be considered highly susceptible to OROV-IRCCS, and thus represent a robust small-animal infection model to assess interventions against OROV.

### 4’-FlU protects AG129 mice and Syrian hamsters against OROV-IRCCS infection, also with delayed treatment

To assess the antiviral activity of 4’-FlU against OROV-IRCCS in AG129 mice, animals were randomly assigned into five groups (n = 5 per group) and administered orally once daily, starting at 2h before infection and then for 7 consecutive days, with either vehicle or 4’-FlU at doses of 10 mg/kg, 5 mg/kg, 2 mg/kg, or 0.4 mg/kg. (Fig. 3a). The once-daily oral dosing schedule was inspired by the reported pharmacokinetic profile in mice with sustained plasma concentrations for over 12h after dosing^26^ and a favorable tolerability profile in AG129 mice (Supplementary Fig. 4). Mice in the vehicle group exhibited progressive weight loss, a decline in surface body temperature and clinical signs including reduced activity and weakness, leading to the onset of HEP beginning as soon as 3 dpi. All mice within this group were euthanized or dead by 6 dpi (Fig. 3b-c; Supplementary Fig. 5). In contrast, none of the treated mice (regardless of the dose regimen) showed clinical evidence of disease, except for some weight loss in the 2 mg/kg and 0.4 mg/kg groups (Fig. 3b-c; Supplementary Fig. 5). At the endpoint (7 dpi), viral RNA in serum and organs from 4’-FlU–treated mice was generally undetectable or remained below the lower limit of quantification (LLOQ) of the RT-qPCR assay (Fig. 3d). In the brain, viral RNA levels were typically reduced by 5 to 6 log10 while in some animals low levels of viral RNA were still detectable. In lung, liver and spleen, even at a dose as low as 0.4 mg/kg, viral RNA levels were reduced by 6 to 9 log_10_. Notably, no infectious virus could be recovered from serum or organs of any treated mouse, regardless of the dose (Fig. 3e), suggesting that the residual viral RNA detected in some of the treated mice likely represents non-infectious or degraded material.

**Figure 3.**
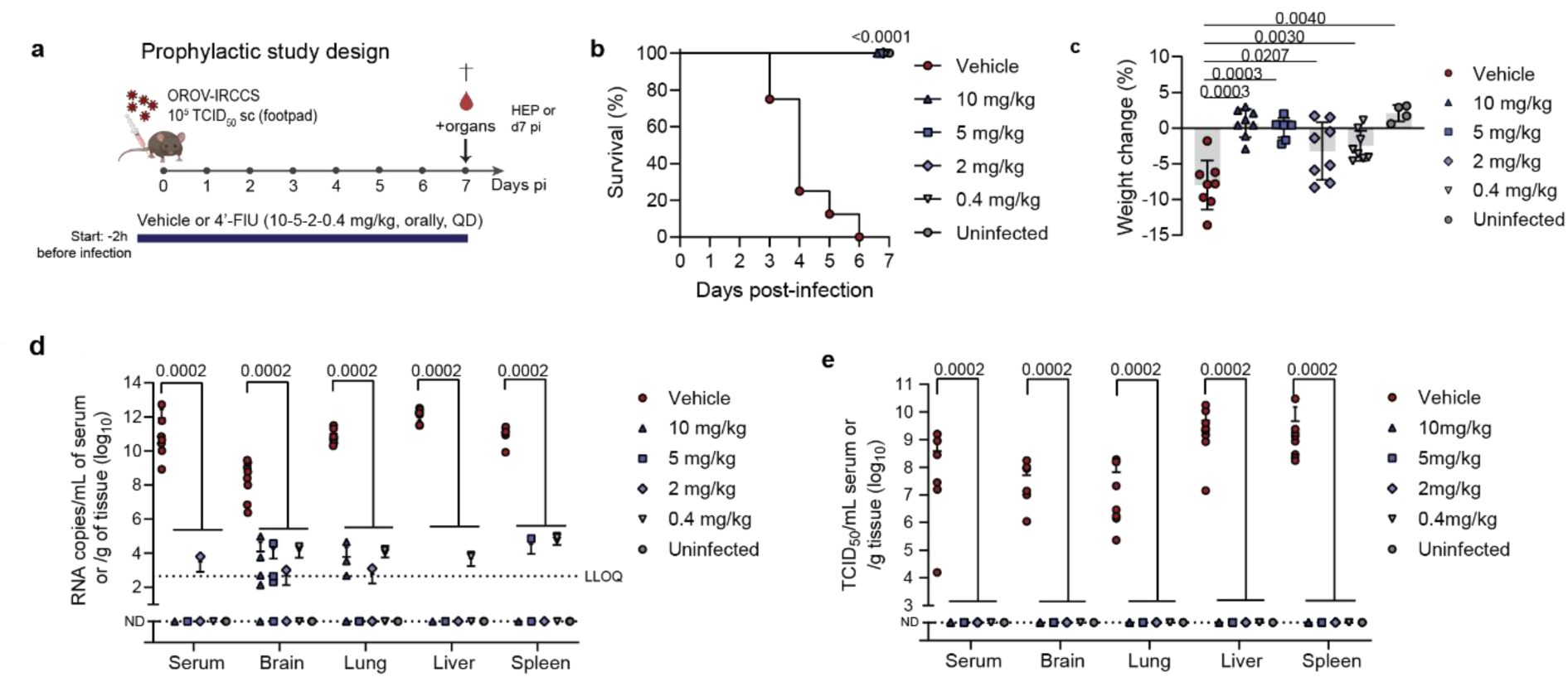
4’-FlU protects against OROV-IRCCS infection in AG129 mice when dosed prophylactically. **(a)** Experimental design scheme. AG129 mice (n=8 per group) were infected subcutaneously in the left rear footpad with 10^5^ TCID_50_ of epidemic OROV-IRCCS strain and treated orally once daily with 4’-FlU (doses ranging from 10 mg/kg to 0.4 mg/kg) starting from 2h prior infection, and during 7 days. Mice were closely monitored until the study endpoint (7 dpi) or until reaching HEP according to a clinical scoresheet. Uninfected mice (n=4) were included as controls. Data presented in this figure have been obtained from 2 biologically independent experiments. **(b)** Kaplan-Meier survival curves display the percentage of survival for each experimental group. Significance was calculated by log-rank (Mantel-Cox) test. **(c)** Weight change of the different experimental groups at time of euthanasia as a percentage, normalized to the body weight at the time of infection. Datapoints are mean ± SD. Significance was calculated by individual Mann-Whitney tests. **(d and e)** Viral RNA loads (d) or infectious titers (e) in serum and various organs measured at the time of euthanasia (HEP or study endpoint) by RT-qPCR and cytopathic effects observation, respectively. Individual animals are presented as dots and error bars represent the SD. Significance was calculated by individual Mann-Whitney tests for non-normally distributed data and by Welch tests for normally distributed data (viral RNA-lung-0.4 mg/kg). Only statistically significant differences are shown. ND, not determined; LLOQ, lower limit of quantification.

Histological analysis was performed on brain, lung, and liver sections collected at the time of euthanasia. In the infected vehicle-treated mice, clear pathological changes were noted, including intra-alveolar hemorrhage in the lungs and extensive coagulative necrosis in the liver (Fig. 4a). No pathological changes were detected in the brain at the HE-staining level (Supplementary Fig. 6). In treated mice, lung and liver architecture appeared preserved across all dose groups. Gross examination of liver tissue from mice in the vehicle group animals revealed pronounced dark coloration, consistent with extensive necrosis. By contrast, livers from treated mice, regardless of the dose, appeared normal and comparable to those of uninfected controls (Supplementary Fig. 7a). Histopathological findings were corroborated by RNAscope *in situ* hybridization (Fig. 4b) The RNAscope probe was custom-designed to target the genomic strand of the viral nucleoprotein (NP) gene. The confocal images revealed abundant viral RNA localized within the brain, lung, and liver of mice in the vehicle group. In contrast, no viral RNA was detected in any organ of mice in the treated groups (Fig. 4b; Supplementary Fig. 8).

**Figure 4.**
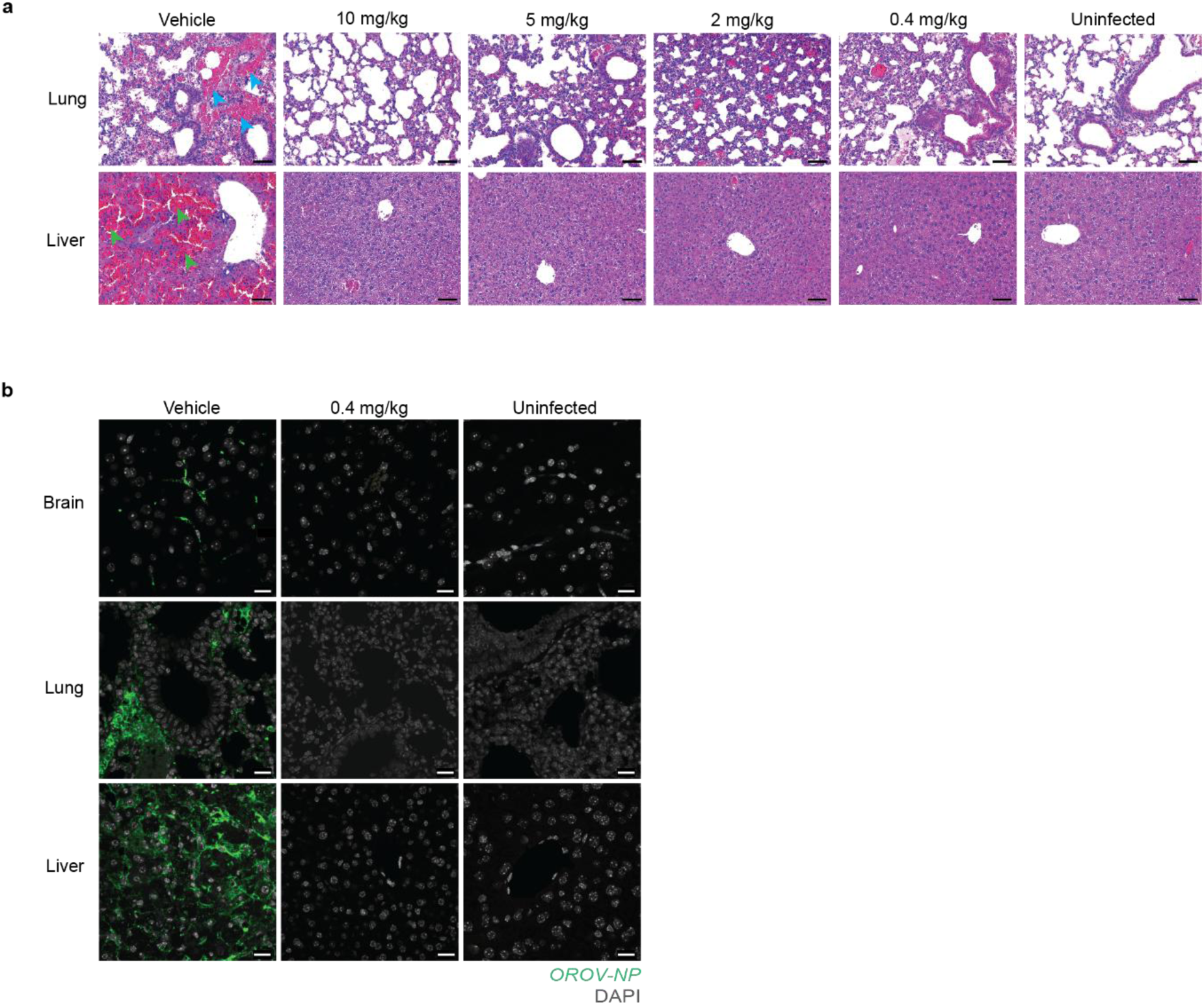
4’-FlU protects against OROV-induced disease when dosed prophylactically in AG129 mice. **(a)** Representative hematoxylin and eosin-stained images of mice lungs and livers obtained at the time of euthanasia (HEP or study endpoint). Blue and green arrowheads indicate intra-alveolar hemorrhage (lungs) and coagulative necrosis (liver), respectively. Scale bar = 50 µm. **(b)** Representative confocal images of 5 µm sections through brains, lungs and liver samples, processed for *in situ* hybridization using the RNAscope platform, with a probe for the OROV S segment, targeting the NP gene. DAPI serves as a nuclear stain. Scale bar = 20 µm.

Next, the antiviral efficacy of 4’-FlU (10 mg/kg, orally, once daily) in OROV-IRCCS infected AG129 mice was also assessed in a therapeutic set-up (Fig. 5a). The start of treatment was delayed until 24, 48, or 72h post-infection (hpi) and continued daily until the study endpoint at 7 dpi (Fig. 5a). All mice in the vehicle group reached the HEP by 5 dpi (Fig. 5b). In contrast, all treated mice, regardless of the timepoint at which treatment was initiated, survived until the study endpoint (Fig. 5b). All mice, also those in which treatment was initiated as late as 72 hpi, remained clinically healthy throughout the study, maintaining body weight and normal surface temperature without observable signs of disease (Fig. 5c, Supplementary Fig. 9). In mice in which treatment was initiated at 24 or 48 hpi, viral RNA levels in serum and organs were mostly either undetectable or below the lower limit of quantification (LLOQ). Even when treatment started as late as 72 hpi, viral RNA levels at endpoint were 1 log_10_ (brain) to 7 log_10_ (liver) lower than in the vehicle group and no infectious virus was recovered from serum or organs (Fig. 5e). In contrast to the mice in the vehicle group, lung and liver architecture remained preserved across all treatment groups (Fig. 7a). Gross examination showed marked dark discoloration of livers from vehicle-treated mice (extensive necrosis), whereas livers from treated mice, irrespective of treatment initiation time, appeared normal and comparable to those of uninfected controls (Supplementary Fig. 7b). Confocal imaging after RNAscope *in situ* hybridization, revealed abundant viral RNA localized within the brain, lung and liver of mice in the vehicle group, but no viral RNA in organs from mice in treated groups, with the exception of a sparse and punctate signal observed in the liver of mice in the 72 hpi group (Fig. 6b; Supplementary Fig. 10). Residual foci of genomic viral RNA, not associated to infectious material, likely persist in the liver of these mice.

**Figure 5.**
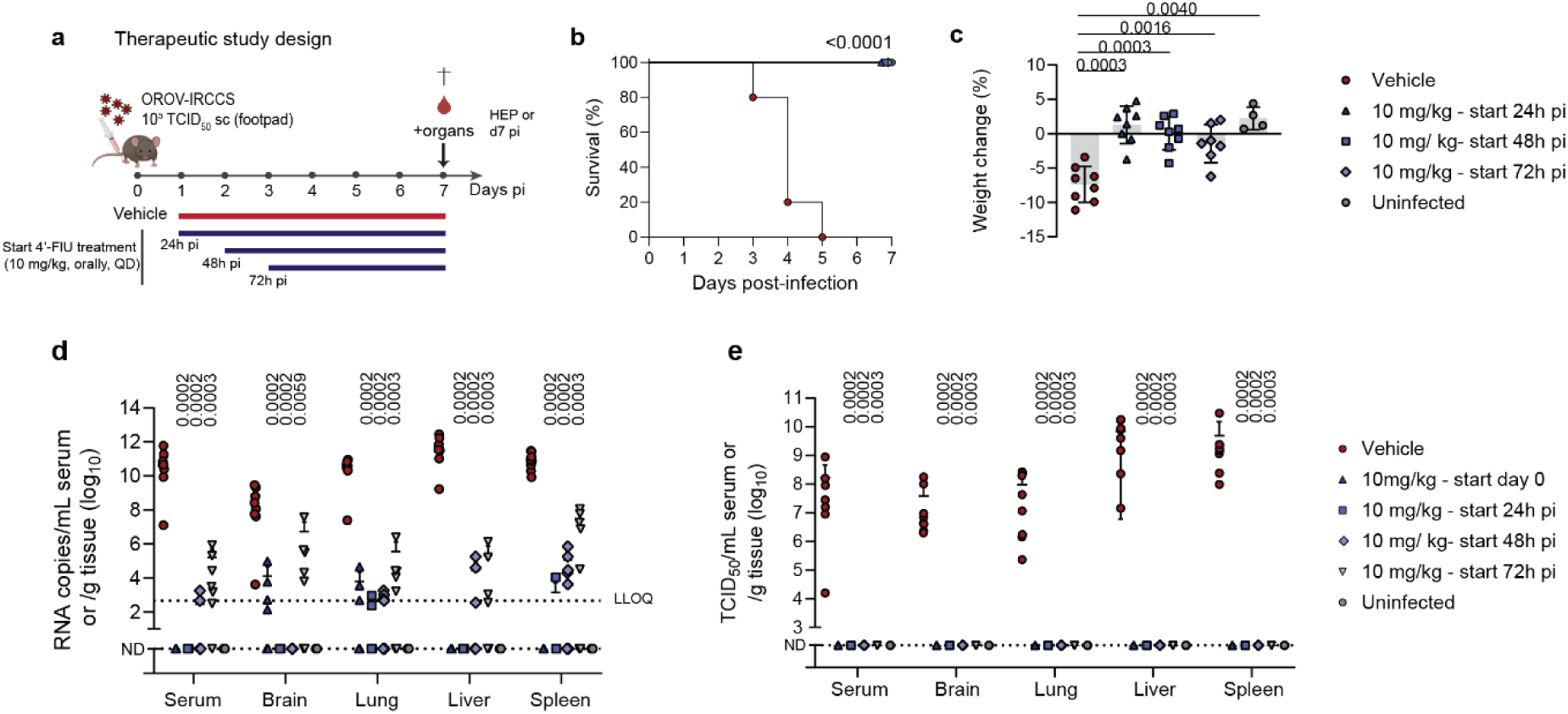
Therapeutic efficacy of 4’FlU against OROV-IRCCS in AG129 mice. **(a)** Experimental design scheme. AG129 mice (n=8 per group, except for the 72h group where n=7 due to one mouse reaching HEP prior to treatment initiation and exclusion from the study) were infected subcutaneously in the left rear footpad with 10^5^ TCID_50_ of epidemic OROV-IRCCS strain and treated orally once daily with 4’-FlU (10 mg/kg) starting either at 24h, 48h or 72h post-infection and during 7 days. Mice were closely monitored until the study endpoint (7 dpi) or until reaching HEP according to a clinical scoresheet. Data presented in this figure have been obtained from 2 biologically independent experiments. **(b)** Kaplan-Meier survival curves display the percentage of survival for each experimental group. Significance was calculated by log-rank (Mantel-Cox) test. **(c)** Weight change of the different experimental groups at time of euthanasia as a percentage, normalized to the body weight at the time of infection. Data points are mean ± SD. Significance was calculated by individual Mann-Whitney tests. **(d** and **e)** Viral RNA loads (d) or infectious titers (e) in serum and various organs measured at the time of euthanasia (HEP or study endpoint) by RT-qPCR and cytopathic effects observation, respectively. Individual animals are presented as dots and error bars represent the SD. The red triangle dataset corresponds to the prophylactic 10mg/kg dataset from Figure 2 and is shown here only for visual comparison. Significance was calculated by individual Mann-Whitney tests for non-normally distributed data and by Welch tests for normally distributed data (viral RNA-spleen-72hpi). Only statistically significant differences are shown. ND, not determined; LLOQ, lower limit of quantification.

**Figure 6.**
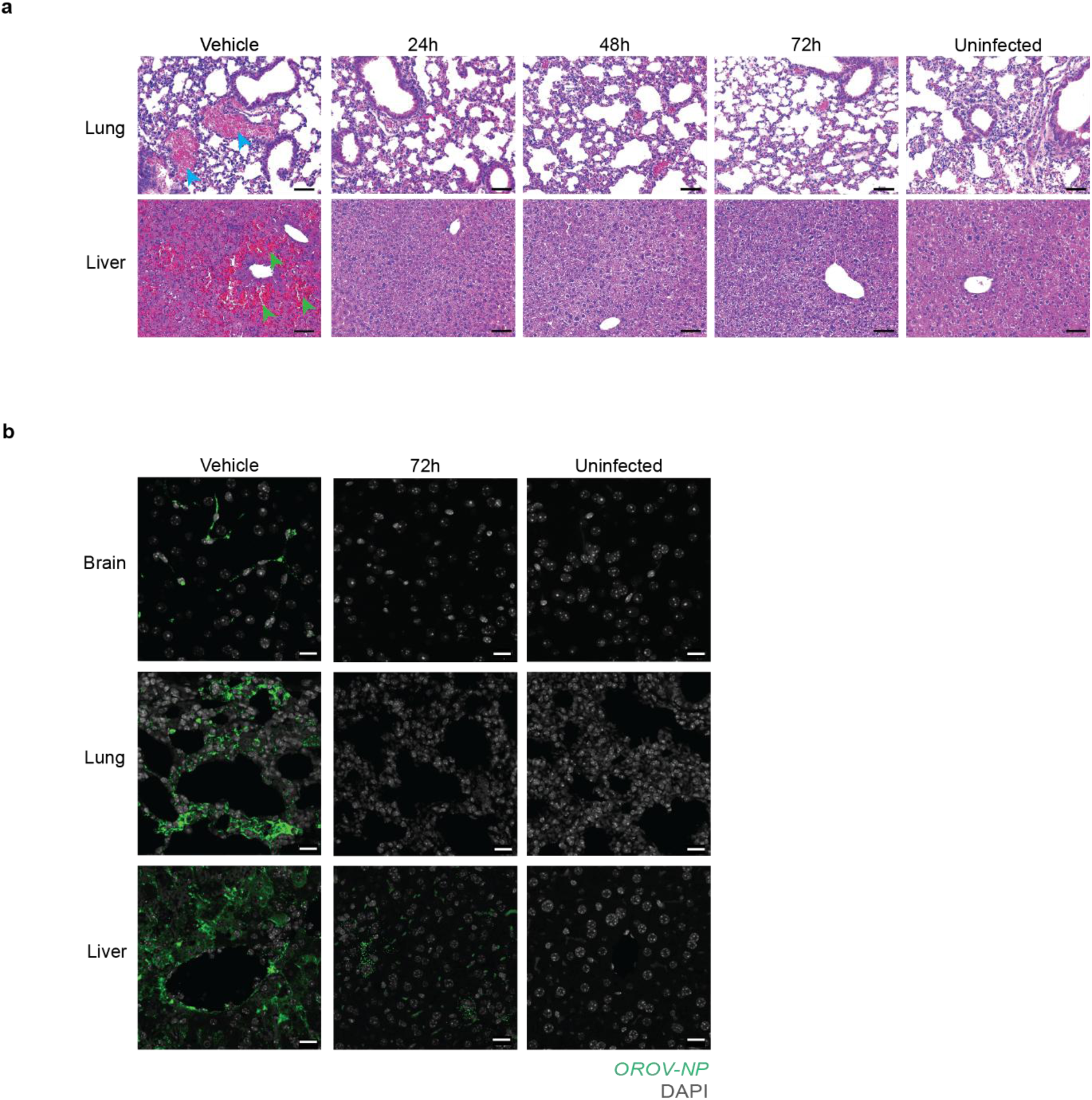
4’-FlU protects against OROV-induced disease when dosed therapeutically in AG129 mice. **(a)** Representative hematoxylin and eosin-stained images of mice lungs and livers obtained at the time of euthanasia (HEP or study endpoint). Blue and green arrowheads indicate intra-alveolar hemorrhage (lungs) and coagulative necrosis (liver), respectively. Scale bar = 50 µm. **(b)** Representative confocal images of 5 µm sections through brains, lungs and liver samples, processed for *in situ* hybridization using the RNAscope platform, with a probe for the OROV S segment, targeting the NP gene. DAPI serves as a nuclear stain. Scale bar = 20 µm.

**Figure 7.**
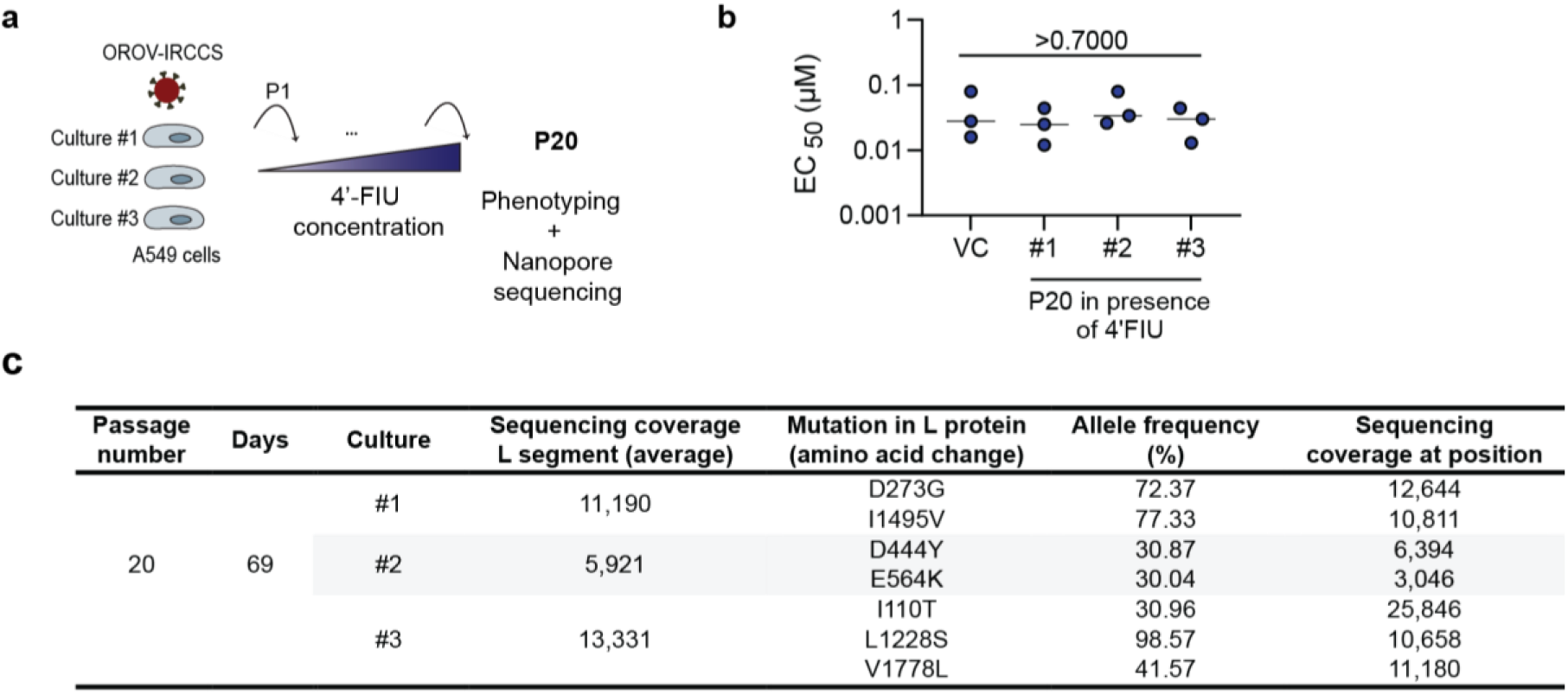
4’-FlU-selected OROV-IRCCS variants are not associated with phenotypic resistance. **(a)** OROV-IRCCS was consecutively passaged in presence of 4’-FlU (three independent cultures), for 69 days (20 passages), in A549 cells. At passage 20 (P20), resulting viral populations were subjected to antiviral assays for phenotyping and sequenced using our in-house Nanopore sequencing pipeline. **(b)** Viral populations obtained under 4’-FlU selective pressure at P20, were subjected to antiviral dose-response assays in A549 cells to determine EC_50_ values. Virus culture passaged in parallel and in absence of 4’-FlU served as control (VC). Data points are mean ± SD of n=3 biologically independent experiments. Significance was calculated by individual Mann-Whitney tests. **(c)** Viral populations obtained under 4’-FlU selective pressure at P20, were subjected to sequencing using our in-house Nanopore pipeline. For each culture, the average L segment sequencing coverage is depicted, and for each identified substitution, the corresponding amino acid change, allele frequency, and coverage at the mutation site are shown.

The antiviral activity of 4’-FlU was finally studied in wild-type Syrian hamsters infected subcutaneously with 10^5^ TCID_50_ of OROV-IRCCS. Animals received 4’-FlU at 10/mg/kg/day in either a prophylactic or therapeutic regimen (Supplementary Fig. 11a). Vehicle-treated hamsters exhibited clinical signs including limb paralysis, weight loss, reduced activity and weakness, leading to the onset of humane endpoints at 3 dpi. All hamsters within this group were euthanized by 5dpi (Supplementary Fig. 11b). Notably, all the treated hamsters reached study endpoint (7dpi) and none showed clinical evidence of disease, but instead gained weight (Supplementary Fig. 11c). Strikingly, no infectious virus could be recovered from any serum or organs of any of the treated hamsters, while animals that received the vehicle showed average values of 7.90×10^8^ TCID_50_/mL serum, 4.92×10^7^ TCID_50_/g brain, 2.60×10^8^ TCID_50_/g lung, 2.02×10^9^ TCID_50_/g liver or 8.74×10^7^ TCID_50_/g spleen (Supplementary Fig. 11d). Histopathological analysis performed at the time of euthanasia revealed marked tissue damage in vehicle-treated hamsters, intra-alveolar hemorrhage in the lungs, and extensive coagulative necrosis in the liver (Supplementary Fig. 11e). However, no histological lesions were observed in the brain, at the HE-staining level. In lungs and liver, 4′-FlU treatment preserved the histological architecture (Supplementary Fig. 11e).

Altogether, the convergent results obtained in AG129 mice and Syrian hamsters, demonstrate that oral administration of 4’-FlU completely blocked viral replication and conferred protection against virus-induced disease, even when dosing was initiated as late as 72 hpi.

### 4’-FlU treatment is characterized by a high barrier to the emergence of resistance

In its 5’-triphosphate form, 4’-FlU has been reported to target the viral polymerase of phylogenetically divergent RNA viruses^24–26^. Ideally, an antiviral drug should have a high barrier-to-resistance. Therefore, we attempted to select for 4’-FlU resistant OROV variant(s) *in vitro,* in three independent cultures, by serially passaging OROV-IRCCS in the presence of suboptimal concentrations of the drug (Fig. 7). After 69 days of passaging (passage 20), the virus populations obtained were independently evaluated for susceptibility to 4’-FlU in antiviral assays (Fig. 7b). Notably, none of the three independent cultures exhibited a measurable shift in EC_50_ value relative to the virus control (VC; passaged in absence of compound) (Fig. 7b). Whole viral genome sequencing was performed using an in-house Nanopore sequencing pipeline, and revealed some substitutions in the viral genome of the three cultures (Fig. 7c), that were not identified in the virus control culture (passaged in absence of 4’-FlU). The mutations were distinct between cultures and occurred at variable allele frequencies, with no convergent substitutions observed (Fig. 7c). Collectively, these findings indicate that prolonged passaging in the presence of 4′-FlU does not select for variants associated with a detectable resistance phenotype, supporting a high genetic barrier to the emergence of resistance.

## DISCUSSION

In an effort to identify potential antiviral drugs that can be repurposed to treat OROV infections, we evaluated a selection of molecules with reported activity against other RNA viruses (obeldesivir, galidesivir, GS-441524, sofosbuvir, ribavirin, favipiravir and molnupiravir). Since OROV encodes, alike other segmented RNA viruses including influenza viruses, an endonuclease, we explored also whether baloxavir, a molecule with reported modest antiviral activity against Bunyamwera virus^21^, can inhibit the replication of OROV. Only 4’FlU resulted in noticeable activity against *in vitro* viral replication. 4’-FlU (also known as EIDD-2749) is a potent broader-spectrum ribonucleoside analog in phase 1 studies, with reported antiviral activity against several RNA viruses including RSV, SARS-CoV-2, IAV, LASV, NiV and CHIKV^23–26^.

To assess whether the potent *in vitro* activity against OROV also translated in protection in an animal infection model, we developed a robust mouse infection model. The interferon (IFN) response has previously been identified as a key determinant of host control over OROV replication^31,32^. AG129 mice, which carry knockouts of type I and II IFN receptor genes, are highly susceptible to multiple viral infections^33–35^. We here observed that these mice are highly susceptible to the epidemic strain OROV-IRCCS. Infected mice developed a severe acute disease that progressed rapidly, characterized by reduced activity, weakness, progressive weight loss and a decline in surface body temperature, with some mice reaching HEP as early as 3 dpi. Prophylactic administration of 4’-FlU, even at a very low dose (0.4 mg/kg/day), conferred complete protection against virus-induced disease, accompanied by preservation of tissue architecture and clearance of infectious virus. The potent antiviral activity was corroborated by a reduction of viral RNA synthesis in organs, as shown by RNAscope *in situ* hybridization staining. Remarkably, such observations were also made when the treatment was initiated as late as 72 hpi, a time at which several infected untreated mice already showed signs of disease. Notably, one mouse in the study already reached HEP at 3 dpi. Thus, 4’-FlU is sufficiently potent to curb an aggressive ongoing infection.

While acknowledging the limitations of the AG129 mouse model, which primarily stem from its profoundly immunocompromised status and the limited neurotropism of OROV in this model, where viral replication is confined to the vasculature, we sought to further validate our findings in the wild-type Syrian hamster model. OROV-IRCCS infection in Syrian hamsters resulted in rapid progression to humane endpoints accompanied by neurological manifestations in 50% of the animals in the vehicle group, thereby providing a more relevant framework to assess the impact of antiviral therapy on OROV brain tropism. Notably, 4′-FlU treatment yielded outcomes consistent with those observed in AG129 mice, with both prophylactic and therapeutic administration protecting animals from disease progression. Infectious virus was undetectable in serum and in all examined organs at study endpoint. These findings demonstrate that the neurological manifestations associated with OROV infection in the hamster model are effectively prevented/treated by 4′-FlU administration.

Antiviral drugs have ideally a high genetic barrier to resistance. We passaged OROV-IRCCS for 69 consecutive days (20 passages) *in vitro* in the presence of suboptimal concentrations of 4’-FlU. None of the resulting three independent viral populations had reduced sensitivity to 4’-FlU. However, some substitutions in the viral genome (L segment) were identified by sequencing, which were distinct between cultures and occurred at variable allele frequencies, with no convergent substitutions observed among cultures. Nevertheless, these mutations did not affect the *in vitro* activity of 4’-FlU. Also the fact that complete clearance of infectious virus could be achieved in two stringent animal models, further points to the high barrier to resistance of this drug. The high genetic barrier to resistance observed in the present study is consistent with previous findings reported for another bunyavirus, Junín virus^25^.

Structural predictive modeling of the OROV L protein, closely aligning with the experimental structure of LACV L protein (Supplementary Fig. 12a-c), revealed a conserved binding pocket accommodating 4’-FlU-TP within the active site, rationalizing its selective incorporation and antiviral mechanism (Supplementary Fig. 12d). Residues forming contacts with the fluorine moiety of 4′-FlU-TP, are conserved in bunyaviruses and across a broader range of negative-sense RNA viruses, including influenza viruses^26^. This conservation supports the extended antiviral activity of 4′-FlU, well beyond the bunyavirus group. Mechanistically, we here demonstrate that 4′-FlU-TP functions as a chain terminator, capable of causing premature termination of peribunyaviru*s* RNA synthesis (despite the presence of a 3’-hydroxyl group) once incorporated into the nascent strand. Given the mechanistic diversity in genome replication and transcription strategies both (i) amongst bunyaviruses on the one hand and (ii) between bunyaviruses and other negative-sense RNA viruses^36^ on the other hand, the potent antiviral activity of 4′-FlU observed in this study may reflect its ability to disrupt multiple critical steps in OROV/bunyavirus genome replication and/or transcription.

Several limitations should be considered when interpreting our findings. First, the absence of an experimentally determined OROV L protein structure required us to infer structural determinants largely through comparison with the related LACV L protein. Although the prediction models generated in this study exhibited high confidence scores and favorable structural quality metrics, the lack of direct structural information for OROV limits definitive mechanistic conclusions and interpretations. Similarly, our *in vitro* replication assays were performed using the LACV system as a surrogate. The inhibitory effects of 4′-FlU on viral replication are consistent with those previously reported for the phylogenetically distant RSV^24^, supporting a conserved mechanism of antiviral action across divergent RNA viruses. An additional consideration relates to the potential safety of 4′-FlU during pregnancy. To our knowledge, the teratogenic potential of 4′-FlU has not yet been formally evaluated. This uncertainty is particularly relevant in the context of OROV infection, as vertical transmission has been clinically documented and associated with adverse fetal outcomes, including congenital disease and miscarriages.

This study identifies 4’-FlU as a promising therapeutic candidate for the treatment of OROV infections. Beyond its efficacy against OROV, our findings also further support the continued development of 4′-FlU as a broader-spectrum antiviral, given its previously demonstrated activity against a range of bunyaviruses, including LASV, Crimean-Congo hemorrhagic fever virus (CCHFV), and Andes virus (ANDV)^25,37^, as well as viruses from more phylogenetically divergent families^23,24,26^. Finally, our study highlights the strategic value of targeting conserved viral structural elements for the development of robust and broad(er) active antiviral agents.

## METHODS

### Cells and viruses

VeroE6 cells (Rhesus monkey kidney, ATCC, CRL-1586) and A549 cells (human lung adenocarcinoma, ATCC, CCL-185) were cultured in Dulbecco’s modified Eagle medium (DMEM, Gibco) supplemented with 10% heat-inactivated fetal bovine serum (FBS, Hyclone) and 1% Non-Essential Amino Acids (NEAA, Gibco). JEG-3 cells (human choriocarcinoma, ATCC, HTB-36) were maintained in Minimum Essential Medium (MEM) supplemented with 10% FBS, 1% sodium pyruvate (Gibco), 1% HEPES (Gibco), 1% NEAA (Gibco) and 1% L-glutamine (Gibco). VeroE6-H2B-miRFP670 cells constitutively expressing near infrared fluorescent protein miRFP-670 were generated by lentiviral transduction and monoclonal selection as described in ^38^ and were maintained in DMEM. Endpoint titrations were performed on VeroE6 cells using medium containing 2% fetal bovine serum, 1% NEAA, 100 U/mL penicillin-streptomycin and 2.50µg/mL amphotericin B (Gibco). For the antiviral experiments (OROV and LACV) with VeroE6, VeroE6-H2B-miRFP670, A549 and JEG3 cells, the respective cell culture medium was used but with 2% FBS instead of 10% FBS. Cell culture incubations were done in humidified incubators at 37°C and 5% CO_2_. Cell lines tested negative for mycoplasma contamination.

The epidemic OROV-IRCCS strain, designated OROV_IRCCS-SCDC_1/2024_, was isolated from a serum sample of a patient returning from Cuba and admitted to the IRCCS Sacro Cuore Don Calabria Hospital in Negrar di Valpolicella (Italy) in June 2024^22^. The whole genome sequence is available in GenBank (BankIt2843215; S segment: PP952117, M segment: PP952118, L segment: PP952119)^5^. The isolate was propagated for 2-3 passages on VeroE6 cells to generate the virus stock used for inoculation, which has a titer of 2.81×10^7^ TCID_50_/mL. The pre-epidemic OROV-TR9760 strain, designated OROV-IP/OROV/TR9760/12/08/2009, was obtained from EVAg (Ref. 014V-02948), and passaged on VeroE6 cells for virus stock production. The LACV/Human/78V strain, designated LACV, was obtained from BEI Resources (Ref. NR-51644), and passaged on VeroE6 cells for virus stock preparation.

### Molecules

For *in vitro* studies, favipiravir, ribavirin and remdesivir were purchased from Carbosynth. GS-441524, molnupiravir, sofosbuvir and obeldesivir were purchased from Excenen Pharmatech Co. Baloxavir marboxil, galidesivir and 4’-fluorouridine were purchased from MedChemExpress.

For animal studies, 4’-Fluorouridine was purchased from Excenen Pharmatech Co. and formulated as follows: 0.5% Tween80 (Sigma) in 10 mM sodium citrate (Sigma) dissolved in water. Compound solutions were prepared daily, one to two hours prior to dosing.

### In vitro antiviral and cytotoxicity assays

VeroE6 or VeroE6-H2B-miRFP670 (25,000 cells/well), A549 (15,000 cells/well) and JEG-3 cells (8,000 cells/well) were seeded in 96-well plates one day before compound treatment and virus infection. The next day (day 0), cells were treated with three-fold serial dilutions of the compounds and infected with OROV at a multiplicity of infection (MOI) of 0.006 (A549 and VeroE6), 0.15 (JEG-3) median TCID_50_ per cell or with LACV at a MOI of 0.01. Three (OROV) or four (LACV) days post infection, the cell viability was determined using staining with 3-(4,5-dimethylthiazol-2-yl)-5-(3-carboxymethoxyphenyl)-2-(4-sulfophenyl)-2H-tetrazolium (CellTiter 96^®^ Aqueous MTS reagent, Ref. G1111, Promega). The percentage of antiviral activity was calculated by subtracting the background and normalizing to the untreated − uninfected control wells, and the EC_50_ was determined using logarithmic interpolation. The potential toxicity of compounds was assessed in a similar set-up in treated − uninfected cultures, in which metabolic activity was quantified at day 3 or day 4 using the MTS assay. The 50% cytotoxic concentration (CC_50_, the concentration at which cell viability reduced to 50%) was calculated by logarithmic interpolation.

### Virucidal and time-of-addition assays

For virucidal assays, a total of 2×10^7^ TCID_50_ of OROV-IRCCS was incubated with fresh medium containing either 1µM 4’-FlU or an equivalent volume of DMSO for 1h at 37°C. Following this, the virus suspension was diluted 1,000 times in fresh medium and used to infect naive A549 cells (50,000 cells/well of a 48-well plate) for 2h at 37°C. After this incubation, the inoculum was removed, the cells were washed 3x with PBS and 500µL of fresh medium was added to the cells. Cells were incubated at 37°C for 24h prior to viral RNA extraction from culture supernatant using the Nucleospin^®^ RNA virus kit (Macherey-Nagel) according to the manufacturer’s instructions. RT-qPCR was conducted as described below and results expressed as RNA copies/mL of culture supernatant.

For time-of-addition assays, A549 cells (15,000 cells/well) were seeded one day prior to infection in 96-well plates. Infection was performed with 1.1×10^7^ TCID_50_ per well, and 1µM 4’-FlU (or the corresponding volume of DMSO) was added to the cells, at the corresponding timepoints : 0h (compound added at the time of infection), 2h, 4h or 7h post-infection. At 10hpi, culture supernatants were removed and cells were washed 3x with PBS prior to intracellular RNA extraction using the Cells-to-cDNA II kit (Invitrogen, Ref. AM1723) according to the manufacturer instructions. RT-qPCR was conducted as described below and results were expressed as relative intracellular RNA levels (% of the corresponding DMSO control).

### Expression and purification of recombinant LACV L_H34K_

The LACV L_H34K_ (strain mosquito/78) construct used for expression encodes full-length LACV L with (i) a H34K mutation that inactivates the ENDO activity^39^ and (ii) an internal Strep-Tag II “WSHPQFEK” flanked by a SG/GSG linker between residues G1034 and Y1035 (G1034-SG-StrepTag II-GSG-Y1035) that allows purification while maintaining replication activity compared to the wild type LACV L construct^29^.

LACV L_H34K_ expressing baculoviruses were made using the Bac-to-Bac method (Invitrogen)^40^. For protein expression, High 5 (Hi5) cells at 0.5 × 10^6^ cells/mL concentration were infected by adding 0.1% of baculovirus and collected 72 hours after the day of proliferation arrest.

Hi5 cells were resuspended in lysis buffer (50 mM Tris–HCl pH 8, 500 mM NaCl, 2 mM β-mercaptoethanol (BME), 5% glycerol) completed with cOmplete EDTA-free protease inhibitor cocktail (Roche) and RibonucleaseA (Sigma-Aldrich), and disrupted by sonication during 3 min (10 sec ON, 20 sec OFF, 50% amplitude) on ice. Lysate was clarified by centrifugation at 48,000 × *g* for 45 min at 4 °C and filtered. The soluble fraction was loaded on a StrepTrap HP column (Sigma-Aldrich) pre-equilibrated in lysis buffer and eluted using lysis buffer supplemented by 2 mM d-Desthiobiotin (Sigma-Aldrich). LACV L_H34K_ fractions were subsequently pooled, dialyzed 1 hour at 4 °C in heparin buffer (50 mM Tris–HCl pH 8, 250 mM NaCl, 2 mM BME, 5% glycerol) and loaded on HiTrap Heparin HP column (Sigma-Aldrich). Elution was performed using 50 mM Tris–HCl pH 8, 1 M NaCl, 5 mM BME, 5% glycerol. The best fractions were flash frozen in liquid nitrogen and stored at −80 °C for future experiments.

### LACV L_H34K_ in vitro replication assay

*In vitro* RNA synthesis was studied by incorporation of radiolabelled ^32^P-GTP in RNA product. For the *de novo* replication assay, 0.57 µM of LACV L_H34K_ was incubated with 0.85 µM of 5’mut_1-19_ (an RNA corresponding to the 19 first nucleotides modified to diminish the complementarity with the 3’ while conserving the hook structure) (5′-ACG AGU GUC GUA CCA AGU A3′) in a buffer containing 50 mM Tris pH 8, 250 mM NaCl and 5 mM BME for 20 min at 4 °C. 0.85 µM of 3′vRNA1–25 (5′-UAU CUA UAC UUG GUA GUA CAC UAC U-3′) was then added for 20 min at 4 °C. Reactions were started by adding 0.1 mM of indicated nucleotides, 56 µCi/µl [^32^P-GTP] and 2 mM MgCl_2_. When indicated, various concentration of 4′-FlU-TP (MedChemExpress) were added prior to the other nucleotides (100µM ATP and GTP) on ice without incubation time. Reactions were incubated at 30 °C for 4 h.

All reactions were stopped by adding 2× volume of loading buffer (95% formamide, 1 mM EDTA, 0.025% SDS, 0.025% bromophenol blue, 0.01% xylene cyanol). The products were denatured by heating to 95 °C for 5 min and loaded on a 20% Tris-Borate-EDTA (TBE)−7M urea-polyacrylamide gel. The RNA products were separated with 1× TBE buffer for 2 h. The gels were exposed on a storage phosphor screen and read with an Amersham Typhoon scanner. The molecular marker is an RNA corresponding to product complementary to the 3′vRNA with a length of 9 (5′-AGU AGU GUA3′) nucleotides with 5′-OH. The MW RNAs were radiolabelled by incubation of the MW RNA at 10 µM with 5 units of T4 polynucleotide kinase (NEB) in its buffer and 0.5 µCi/µl [γ-^32^P] ATP at 37 °C during 10 min. Reactions were stopped by incubation at 70 °C for 10 min and the addition of loading buffer. Quantification of Fig. 2 was performed, based on three gels ran with distinct samples, using ImageJ.

### OROV infection in mice and hamsters

Males and females AG129 mice (129/Sv mice with knockouts in both IFNα/β and IFNγ receptor genes) were bred in house. Breeding pairs of AG129 mice had been purchased from Marshall BioResources. The specific pathogen-free status of the mice was regularly checked at the KU Leuven animal facility of the Rega Institute. Female wild-type Syrian hamsters were purchased from Janvier Laboratories. OROV inoculation of AG129 mice was performed in 6-12 week-old animals and inoculation of Syrian hamsters was performed in 6-8 week-old animals. Sex distribution per experimental group is shown in Supplementary Table 3. Animals were housed in individually ventilated isolator cages (IsoCage N Biocontainment System, Tecniplast) at a temperature of 21°C, humidity of 55%, and subjected to 12:12 dark/light cycles. Animals had access to food and water *ad libitum,* and their cages were enriched with cotton and cardboard play tunnels. Animal work was performed under Biosafety Level 3 conditions. Housing conditions and experimental procedures were approved by the Animal Ethics Committee of KU Leuven (approval numbers 202/2024, 036/2025, 021/2026 and M003/2025) following institutional guidelines approved by the Federation of European Laboratory Animal Science Associations (FELASA).

Mice were anesthetized by inhalation with isoflurane before infection. Virus suspension consisting of 10^5^ TCID_50_, prepared in Opti-MEM medium, was injected subcutaneously in the rear of the left footpad of the mice or between the shoulder blades of Syrian hamsters. At humane endpoint or study endpoint, animals were euthanized by administration of Dolethal before sample collection. Animal body weight, signs of disease and surface body temperature (only for mice studies, using a Gentle Temp 720 device, Omron) were monitored at least twice daily. Animals were euthanized according to the following clinical criteria: huddling (2 points), hunch back, ruffled fur, eye recession, hyperactivity, weight loss 5% of baseline (3 points each), ataxia, circling, weakness, shaking, tremors, anemia, 28°C> surface body temperature <30°C, weight loss 10% of baseline (5 points each) and bleeding, paralysis, moribund, unresponsive, surface body temperature < 28°C, weight 20% of baseline (10 points each). The humane endpoint was defined when the score reached ≥ 10.

### Treatment regiment

In prophylactic setting experiments, AG129 mice were administered orally either vehicle (0.5% Tween80 in 10 mM sodium citrate dissolved in water) (n=8), 10 mg/kg (n=8), 5 mg/kg (n=8), 2 mg/kg (n=8) or 0.4 mg/kg (n=8) 4’-FlU, once daily from two hours prior infection to study endpoint. Uninfected mice were used as controls (n=4).

In therapeutic setting experiments, AG129 mice were administered orally either vehicle (n=8) or 10 mg/kg (n=8) 4-FlU, once daily, 24hpi, 48hpi or 72hpi to study endpoint. In the “72h” group, one animal reached humane endpoint before the initiation of the compound treatment and was therefore excluded from the dataset (n=7 for this group). Uninfected mice were used as controls (n=4).

In the Syrian hamsters study, animals were administered orally either vehicle (n=4) or a fixed dose of 10 mg/kg 4’-FlU, prophylactically (n=6) or therapeutically (n=6).

### Endpoint/HEP virus titration assay

Serum was obtained from total blood by centrifugation at 10.000x g for 5 min. Organs were homogenized using bead disruption in 400 µL of DMEM medium and centrifuged (10.000x g, 5 min) to pellet cell debris. To quantify infectious OROV particles, endpoint titrations were performed on VeroE6 cells (10^4^ cells/well) infected with 10-fold sample serial dilutions, in 96-well plates. After seven days, viral titers were determined by the Reed-Muench method^41^ using the Lindenbach calculator and were expressed as 50% tissue culture infectious dose (TCID_50_) per mL of serum or g of tissue. The limit of quantification (LOQ) of the titration assay was set based on the lowest measured virus titer taking tissue weight into account. For statistical analysis, negative samples were assigned the value of the LOQ (1.78×10^2^ TCID_50_/mL or /g).

### RNA extraction and RT-qPCR

Serum was obtained as described above and RNA was extracted using the Nucleospin^®^ RNA virus kit (Macherey-Nagel) according to the manufacturer’s instructions. Organs were collected and homogenized using bead disruption (Precellys) in 700 µL of TRK lysis buffer (E.Z.N.A.^®^ Total RNA kit, Omega Bio-Tek) and centrifuged (15.000x g, 10 min) to pellet cell debris. RNA was extracted according to the manufacturer’s instructions.

RT-qPCR was conducted on a QuantStudio5 platform (Applied Biosystems) using the iTaq Universal probe One-Step RT-qPCR kit (Ref. 1725141, Bio-Rad). The primers targeting the Gc gene on the genomic M segment of OROV-IRCCS were: forward-5’-ACGCAGACTAACCAACTTCC-3’ and reverse-5’-CAGTTCCGATAGTTGACCCATTA-3’.

The probe was 5’-TGCAGTGCGGCTCAATCCAAGTAA-3’. Primers and probes were purchased from Integrated DNA Technologies, Inc (IDT). Standards of OROV-IRCCS cDNA (IDT) were used to express viral genome copies per mL of serum or mg of tissue. The LLOQ was set based on the lowest measured titers of quantified standards. For statical analysis, samples showing values below the LOD (determined as 9.77 RNA copies/mL), were assigned the LOD value.

### Blood count and serum biochemistry analysis

Complete blood counts were performed on whole blood samples, collected in Tris-EDTA buffer solution (Sigma) using the Element HT5^TM^ (Heska) hematology analyzer.

For serum biochemistry analysis, the serum was isolated from whole blood samples as described above, transferred and analyzed at the clinical laboratory of the University Hospital of Leuven using the Cobas 8000 device (Roche). Albumin quantification was done using Bromocresol green method, AST/AST by colorimetric assay according to the IFCC method, ALP by colorimetric assay with 2-amino-2-methyl-1-propanol buffer, Sodium/potassium/chloride by potentiometry with an indirect ion-selective electrode, creatinine using an enzymatic reaction with creatinine peroxidase and total bilirubin by colorimetric assay using the DPD method.

### Histology

Brain, lungs and liver samples were fixed overnight in 4% paraformaldehyde (ThermoScientific) and embedded in paraffin. Tissue sections (5 µm) were stained with hematoxylin and eosin and analyzed blindly by expert pathologists.

### RNAscope *in situ* hybridization

Probe V-OROV-S-O5-sense-C1 (Advanced Cell Diagnostics, Ref. 1811921-C1) was custom-designed against the S segment nucleoprotein gene, reverse-complement of nt 10-937 of the Genbank PP952117.3 sequence^5^. This probe is publicly available. Manual RNAscope assays were performed on 5 µm FFPE sections of brain, lung and livers using the Multiplex Fluorescent Detection Kit v2 (Advanced Cell Diagnostics, Ref. 323110) according to the manufacturer’s protocol and as previously described^34^. Briefly, slides were baked at 60°C for 1 h, followed by deparaffinization in xylene (Leica Biosystems, Ref. 3803665EG), a 100% ethanol wash, and incubation with hydrogen peroxide for 15 min at room temperature. Tissue pretreatment was performed by steaming in target retrieval reagent (Advanced Cell Diagnostics, Ref. 322000) for 15 min followed by incubation in Protease Plus (Advanced Cell Diagnostics, Ref. 322330) at 40°C for 20 min. Signal amplification was followed by development with the fluorophore dye Opal 520 (Akoya Biosciences, Ref. FP1487001KT). DAPI (Thermo Fisher Scientific, Ref. D1306) serving as nuclear stain. Slides were mounted in Mount Solid antifade (Abberior, Ref. MM-2011-2X15ML). Confocal images were taken on a Zeiss LSM 980.

### Resistance mutation selection and Oxford Nanopore sequencing

OROV-IRCCS was passaged in A549 cells in the presence of increasing concentrations of 4’-FlU. Selection was initiated at 10 nM and, for each passage, the virus was cultured at the same concentration, and at 3×, 10× and 30× higher concentrations. The culture with the highest compound concentration that still showed virus breakthrough, as observed by a significant cytopathic effect, was then used for the next passage. At passages 20 (day 69), vRNA in the cell-culture medium of three independent lineages was extracted as described above or culture supernatant was used in antiviral assays (as described above) to assess 4’-FlU phenotypic susceptibility. Following extraction, purified viral RNA was converted to cDNA by sequence-independent, single-primer amplification as previously described^42^. Amplified cDNA was sequenced on an Oxford Nanopore Technologies GridION device (MinKNOW v25.03.7) using real-time super-accurate basecalling (Dorado v7.8.3). SISPA adapters were removed from the obtained reads using Porechop (https://github.com/rrwick/Porechop), after which variant calling was performed using an in-house script that performs reference mapping with minimap2^43^, followed by BAM file inspection with bam-readcount^44^ to obtain a detailed overview of nucleotide variation at each position. Only sites with >20x coverage were included in the analysis and only variants with >10% prevalence and >10x variant-specific coverage were assessed.

### Structure predictions

The OROV L protein sequence (UniPRot A0A0B5CUJ1) was used together with short RNA sequences as input for either AlphaFold3 (AF3)^45^ through the AF3 server at https://alphafoldserver.com/ or for a local implementation of Boltz-2^46^, for modeling with nucleotide analogues including 4’-FlU-TP, which were input as SMILES strings. Results were visualized and analyzed with Pymol (The PyMOL Molecular Graphics System, Version 3.0 Schrödinger, LLC) and ChimeraX^47^. Comparisons with La Crosse virus polymerase used Protein Data Bank entries 7ORI to 7ORM^48^. In Supplementary Fig. 12, 7ORK, which harbors an ATP at the catalytic site, is used.

### Sequence alignments

Viral L protein sequence alignments were performed using MEGA12^49^. Genbank sequence accession numbers for L segments, from which L protein sequence were deduced, are: PP952119 (OROV-IRCCS), KP026179.1 (OROV-TR9760), AF528165.1 (LACV/Human78V), (LASV-Josiah), AF291704.5 (Andes-Chile9717869), AY422209.2 (CCHFV-IbAr10200).

### Graphs and statistical analyzes

All graphs and statistical analyses were performed using GraphPad Prism version 10.4.1. Normality of the data distribution was assessed using the Shapiro-Wilk test. Survival analyses were conducted using the log-rank (Mantel-Cox) test. For comparisons involving groups or datasets with non-normally distributed data, unpaired Mann-Witney tests (two-sided) were used. For comparisons involving groups or datasets with normally distributed data, Welch tests (two-sided) were used.

## SUPPLEMENTARY INFORMATIONS

Supplementary figures 1 to 12 ; Supplementary tables 1 to 3.

## ACKNOWLEDGEMENTS

We thank Tina Van Buyten for growing the OROV-TR9760 stock and initial antiviral assays. We thank Steven De Jonghe and Wim Smet for compound ordering, Thibault Francken for culturing the VeroE6-H2B-miRFP670 cells, the colleagues at the Rega animal facility at KU Leuven and Madalen Le Gorrec (University Grenoble Alpes) for technical assistance. This work benefited from the platforms of the Grenoble Instruct-ERIC Center (ISBG, UAR 3518 CNRS– CEA–UGA–EMBL), within the Grenoble Partnership for Structural Biology (PSB), supported by FRISBI (ANR-10-INBS-0005). The IBS acknowledges its integration within the Interdisciplinary Research Institute of Grenoble (IRIG, CEA).

This work was funded by European project DURABLE (Delivering a Unified Research Alliance of Biomedical and public health Laboratories against Epidemics, to JN), by the ANRS-Maladies Infectieuses Emergentes (grant ANRS0380 to SB) and by GRAL, a program from the Chemistry-Biology-Health (CBH) Graduate School of University Grenoble Alpes (grant ANR-17-EURE-0003 to HM). The funders had no role in the study design, data collection and interpretation, or the decision to submit the work for publication.

## AUTHORS CONTRIBUTION

MF and MD: Project design, design and execution of the experiments, data analysis and manuscript writing. IT and HM: Design, Execution and analysis of the in vitro replication assays. TB and SB: Design, execution and analysis of the structural prediction work. MK : Design and execution of the RNAscope experiments. PM: Design of RNAscope experiments. TMP and BV: Execution and analysis of the sequencing work. TR, BW and DRT: analysis of histopathology data. NC, SH: Execution of experiments. KD: Execution of in vitro antiviral assays and resistance selection experiments. VL : Execution of sequences alignment analysis. JRP: Access to OROV-TRVL9760 strain. CC: Access to OROV-IRCCS strain. ML, JN: Funding acquisition, project design and manuscript writing. All authors reviewed the manuscript, provided feedback, and approved the final version for submission.

## DATA AND MATERIALS AVAILABILITY

All data generated and analyzed in this paper are included in this article. Additional imaging data related to the RNAscope analysis are available from the corresponding authors upon request. Source data are provided with this paper.

## REFERENCES

1. de Souza, W. M. et al. ICTV Virus Taxonomy Profile: Peribunyaviridae 2024. J. Gen. Virol. 105, 002034 (2024).

2. Anderson, C. R., Spence, L., Downs, W. G. & Aitken, T. H. G. Oropouche Virus: a New Human Disease Agent from Trinidad, West Indies. Am. J. Trop. Med. Hyg. 10, 574– 578 (1961).

3. Bai, F., Denyoh, P. M. D., Urquhart, C., Shrestha, S. & Yee, D. A. A Comprehensive Review of the Neglected and Emerging Oropouche Virus. Viruses 17, 439 (2025).

4. PAHO/WHO. Epidemiological Update Oropouche in the Region of the Americas - 13 August 2025. (2025).

5. Deiana, M. et al. Full Genome Characterization of the First Oropouche Virus Isolate Imported in Europe from Cuba. Viruses 16, 1586 (2024).

6. Migueres, M. et al. Oropouche Fever: An Imported Case Underscoring the Importance of Clinical and Epidemiological Vigilance. Travel Med. Infect. Dis. 102862 (2025).

7. Gourjault, C. et al. Persistence of Oropouche virus in body fluids among imported cases in France, 2024. Lancet Infect. Dis. 25, e64–e65 (2025).

8. Có, A. C. G. et al. Unravelling the pathogenesis of Oropouche virus. Lancet Infect. Dis. 25, e381–e382 (2025).

9. Naveca, F. G. et al. Human outbreaks of a novel reassortant Oropouche virus in the Brazilian Amazon region. Nat. Med. 30, 3509–3521 (2024).

10. Sah, R. et al. Oropouche fever fatalities and vertical transmission in South America: implications of a potential new mode of transmission. Lancet Reg. Health Am. 38, 100896 (2024).

11. PAHO/WHO. Epidemiological Alert in the Region of the Americas: Vertical Transmission under Investigation - 01 August 2024. (2024).

12. Martins-Filho, P. R., Carvalho, T. A. & Dos Santos, C. A. Oropouche fever: reports of vertical transmission and deaths in Brazil. Lancet Infect. Dis. 24, e662–e663 (2024).

13. Ceccarelli, G. et al. Oropouche virus infection: Differential clinical outcomes and emerging global concerns of vertical transmission and fatal cases. Int. J. Infect. Dis. 150, 107295 (2025).

14. Schwartz, D. A., Baud, D. & Dashraath, P. A potential mechanism of transplacental transmission of Oropouche virus in pregnancy. Lancet Microbe 101083 (2025).

15. Giovanetti, M. Oropouche virus and the urgent need for global surveillance. Nat. Microbiol. 10, 2–3 (2025).

16. Elliott, R. M. Orthobunyaviruses: recent genetic and structural insights. Nat. Rev. Microbiol. 12, 673–685 (2014).

17. Gutierrez, B. et al. Evolutionary Dynamics of Oropouche Virus in South America. J. Virol. 94, e01127–19 (2020).

18. Scachetti, G. C. et al. Re-emergence of Oropouche virus between 2023 and 2024 in Brazil: an observational epidemiological study. Lancet Infect. Dis. 25, 166–175 (2025).

19. Livonesi, M. C., De Sousa, R. L. M., Badra, S. J. & Figueiredo, L. T. M. In vitro and in vivo studies of ribavirin action on Brazilian Orthobunyavirus. Am. J. Trop. Med. Hyg. 75, 1011–1016 (2006).

20. Yoosuf, B. T. et al. Epidemiology, transmission dynamics, treatment strategies, and future perspectives on Oropouche virus. Diagn. Microbiol. Infect. Dis. 113, 116882 (2025).

21. Ter Horst, S., et al. Enhanced efficacy of endonuclease inhibitor baloxavir acid against orthobunyaviruses when used in combination with ribavirin. J. Antimicrob. Chemother. 75, 3189–3193 (2020).

22. Castilletti, C. et al. Replication-Competent Oropouche Virus in Semen of Traveler Returning to Italy from Cuba, 2024. Emerg. Infect. Dis. 30, 2684–2686 (2024).

23. Yin, P. et al. 4’-Fluorouridine inhibits alphavirus replication and infection in vitro and in vivo. mBio 15, e0042024 (2024).

24. Sourimant, J. et al. 4’-Fluorouridine is an oral antiviral that blocks respiratory syncytial virus and SARS-CoV-2 replication. Science 375, 161–167 (2022).

25. Welch, S. R. et al. Delayed low-dose oral administration of 4’-fluorouridine inhibits pathogenic arenaviruses in animal models of lethal disease. Sci. Transl. Med. 16, eado7034 (2024).

26. Lieber, C. M. et al. 4’-Fluorouridine mitigates lethal infection with pandemic human and highly pathogenic avian influenza viruses. PLoS Pathog. 19, e1011342 (2023).

27. Reguera, J., Weber, F. & Cusack, S. Bunyaviridae RNA polymerases (L-protein) have an N-terminal, influenza-like endonuclease domain, essential for viral cap-dependent transcription. PLoS Pathog. 6, e1001101 (2010).

28. Arragain, B. et al. Pre-initiation and elongation structures of full-length La Crosse virus polymerase reveal functionally important conformational changes. Nat. Commun. 11, 3590 (2020).

29. Arragain, B. et al. Structural snapshots of La Crosse virus polymerase reveal the mechanisms underlying Peribunyaviridae replication and transcription. Nat. Commun. 13, 902 (2022).

30. de Mendonça, S. F. et al. Evaluation of Aedes aegypti, Aedes albopictus, and Culex quinquefasciatus Mosquitoes Competence to Oropouche virus Infection. Viruses 13, 755 (2021).

31. Proenca-Modena, J. L. et al. Oropouche virus infection and pathogenesis are restricted by MAVS, IRF-3, IRF-7, and type I interferon signaling pathways in nonmyeloid cells. J. Virol. 89, 4720–4737 (2015).

32. Sterling, C. E. et al. Oropouche virus causes acute hepatitis in mice controlled by type I interferons. J. Virol. e0061126 (2026).

33. Kaptein, S. J. F. et al. A pan-serotype dengue virus inhibitor targeting the NS3-NS4B interaction. Nature 598, 504–509 (2021).

34. Lin, Y. et al. A robust mouse model of HPIV-3 infection and efficacy of GS-441524 against virus-induced lung pathology. Nat. Commun. 15, 7765 (2024).

35. Segura Guerrero, N. A., Sharma, S., Neyts, J. & Kaptein, S. J. F. Favipiravir inhibits in vitro Usutu virus replication and delays disease progression in an infection model in mice. Antiviral Res. 160, 137–142 (2018).

36. Malet, H., Williams, H. M., Cusack, S. & Rosenthal, M. The mechanism of genome replication and transcription in bunyaviruses. PLoS Pathog. 19, e1011060 (2023).

37. Shrivastava-Ranjan, P. et al. Potent In Vitro Antiviral Activity of 4’-Fluorouridine Against Diverse Orthohantaviruses including Andes Virus. Antiviral Res. 106467 (2026).

38. Chiu, W. et al. Development of a robust and convenient dual-reporter high-throughput screening assay for SARS-CoV-2 antiviral drug discovery. Antiviral Res. 210, 105506 (2023).

39. Reguera, J., Weber, F. & Cusack, S. Bunyaviridae RNA polymerases (L-protein) have an N-terminal, influenza-like endonuclease domain, essential for viral cap-dependent transcription. PLoS Pathog. 6, e1001101 (2010).

40. Nie, Y., Bellon-Echeverria, I., Trowitzsch, S., Bieniossek, C. & Berger, I. Multiprotein complex production in insect cells by using polyproteins. Methods Mol. Biol. Clifton NJ 1091, 131–141 (2014).

41. Reed, L. J. & Muench, H. A SIMPLE METHOD OF ESTIMATING FIFTY PER CENT ENDPOINTS12. Am. J. Epidemiol. 27, 493–497 (1938).

42. Vanmechelen, B. et al. The characterization of multiple novel paramyxoviruses highlights the diverse nature of the subfamily Orthoparamyxovirinae. Virus Evol. 8, veac061 (2022).

43. Li, H. New strategies to improve minimap2 alignment accuracy. Bioinforma. Oxf. Engl. 37, 4572–4574 (2021).

44. Khanna, A. et al. Bam-readcount - rapid generation of basepair-resolution sequence metrics. J. Open Source Softw. 7, 3722 (2022).

45. Abramson, J. et al. Accurate structure prediction of biomolecular interactions with AlphaFold 3. Nature 630, 493–500 (2024).

46. Passaro, S. et al. Boltz-2: Towards Accurate and Efficient Binding Affinity Prediction. Preprint at 10.1101/2025.06.14.659707 (2025).

47. Meng, E. C. et al. UCSF ChimeraX: Tools for structure building and analysis. Protein Sci. Publ. Protein Soc. 32, e4792 (2023).

48. Arragain, B. et al. Structural snapshots of La Crosse virus polymerase reveal the mechanisms underlying Peribunyaviridae replication and transcription. Nat. Commun. 13, 902 (2022).

49. Kumar, S. et al. MEGA12: Molecular Evolutionary Genetic Analysis Version 12 for Adaptive and Green Computing. Mol. Biol. Evol. 41, msae263 (2024).

